# LFASS: Label-Free Automated Survival Scoring for High-Throughput Nematode Assays

**DOI:** 10.1101/195347

**Authors:** Alexandre Benedetto, Timothee Bambade, Catherine Au, Jennifer M Tullet, Hairuo Dang, Kalina Cetnar, Filipe Cabreiro, David Gems

**Affiliations:** Institute of Healthy Ageing, University College London, London, UK.; Division of Biomedical and Life Sciences, Lancaster University, Lancaster, UK.; School of Biosciences, University of Kent, Canterbury, UK.; Institute of Structural and Molecular Biology, University College London, London, UK.

**Keywords:** ageing, autophagy, blue fluorescence, *C. elegans*, high-throughput, *hsf-1*, infection, LFASS, *skn-1*, stress, survival

## Abstract

*Caenorhabditis elegans* is an excellent model for high-throughput experimental approaches, but lacks a robust, versatile and automated means to pinpoint time of death. Here we describe an automated, label-free, high-throughput method using death-associated fluorescence to monitor nematode survival, which we apply to stress and infection resistance assays. We demonstrate its use to define correlations between age, longevity and stress-resistance, and reveal an autophagy-dependent increase in acute stress resistance in early adulthood.

---

Survival assays are widely performed in both basic and applied biomedical research to assess the frailty of organisms, tissues and cells, and to determine the toxicity or efficacy of a chemical agent. In such assays, death is usually revealed by a light signal and/or an enzymatic reaction, requiring reagents that may interfere with the process under study^1^. The nematode *C. elegans* is a widely used experimental model in basic, pharmacological and environmental research^2^, and for the study of parasitic worm species^3, 4^ that affect crops, cattle, and 3.5 billion people^5^. Its small size, optical transparency, and good genetics make it a convenient model organism for high-throughput chemical and bacterial library screens for the development of anti-helminthic and anti-ageing drugs, and for elucidating the biology of host-pathogen interactions^6^. Since the late 1990s *C. elegans* screening platforms have evolved to include microfluidics and automated robotic arms^7, 8^. Yet full automation of *C. elegans* survival assays as been limited by death scoring techniques. Worms are deemed dead when they fail to move in response to touch, making death scoring by movement-tracking impracticable in quiescent, old or mutant animals with impaired mobility or touch-response, or in assays that affect worms’ ability to sense or move.

We recently discovered that an endogenous burst of blue fluorescence - dubbed “death fluorescence” (DF) - generated by de-quenching of anthranilic acid species, occurs in nematodes’ intestine at the onset of death^9^. From time-lapse recordings of blue fluorescence during killing assays, we observed that the timing of individual DF events follows a normal distribution. Considering the total fluorescence of a worm population over time, we found that its half-maximal fluorescence corresponds to half the worms undergoing DF (**Supplementary Figure 1a, 1b**). Hence, the time of half-maximal fluorescence corresponds to the median time of death, which is the main parameter sought in survival assays. Because it is endogenous and easily detectable by fluorescence microscopes, spectrophotometers and plate-readers, DF recording could replace vital dyes and movement tracking algorithms in survival assays in a variety of laboratory setups.

We used DF measurement to develop new, label-free, high-throughput and automated oxidative-stress, heat-shock and infection survival assays. To enable multiple conditions to be tested in parallel and to limit phototoxicity (data not shown), we opted for a 384-well plate format to be read by a light-emitting diode (LED)- illumination plate-reader (**Fig. 1a**). In these conditions, DF was optimally recorded with excitation/emission wavelengths of 360/435 nm (**Supplementary Figure 1c**) using as few as 16 worms per well. With this setup, each measurement takes 0.8 s, allowing for 150 samples to be measured in under 2 min. This enables accurate and unbiased measurement of median time of death in assays spanning from hours (acute stress assays, **Fig. 1b, 1c**) to days (bacterial infection assays, **Fig. 1d**). Hence this label-free automated survival scoring method (henceforth referred to as LFASS) may be adapted for high-throughput approaches.

**Figure 1.**
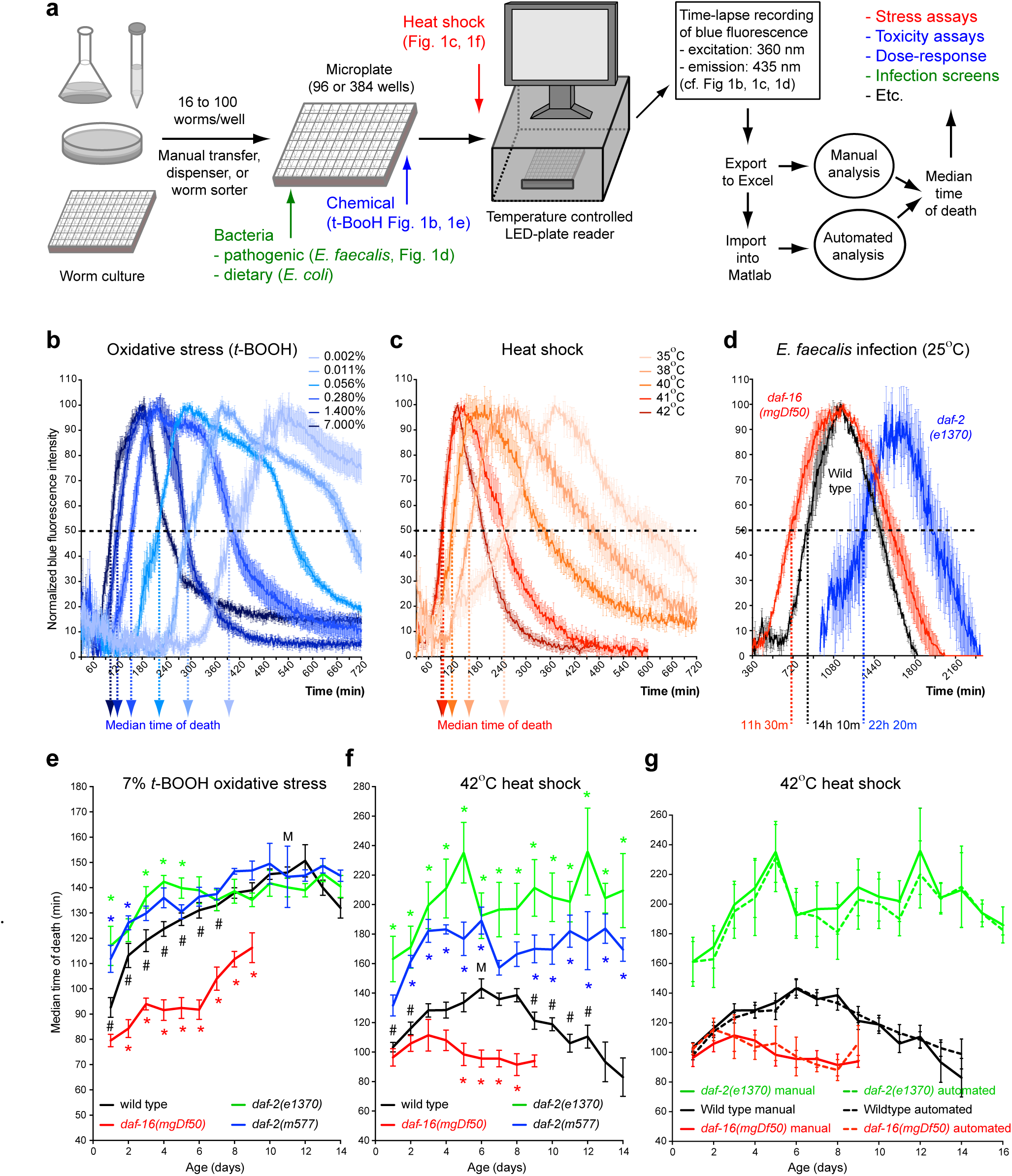
LFASS provides robust automated scoring and analysis of *C. elegans* survival assays. (a) The LFASS pipeline. Dose-dependency of *t*-BOOH-induced oxidative stress (b) and heat-shock (c) resistance measured by LFASS.(d) Automated measurement of resistance to *E. faecalis* bacterial infection by LFASS discriminates between infection sensitive and resistant IIS mutants. LFASS reveals distinct patterns of age-dependent acute stress resistance upon oxidative stress (e) *vs* heat-shock (f) in *C. elegans* IIS mutants. (g) LFASS automated data analysis package and manual analysis yield near-identical results. Error bars, SEM. M: peak resistance for wild-type. Comparison to age-matched wild-type: * *p*<0.05 down to *p*<0.0001. Comparison to M within wild-type values # p<0.05 down to *p*<0.0001

To demonstrate its practicality, we used LFASS to re-examine in unprecedented depth the controversial link between stress resistance and longevity^10-12^, which is tedious to study by traditional means. Survival stress assays have long been used to dissect mechanisms of disease and ageing, revealing that animals lose their ability to endure chronic stress as they age^13-15^, in line with molecular damage theories of ageing^16-18^. However, this correlation has not been rigorously tested with respect to intense, acute stress paradigms that challenge initial resistance to stress, as opposed to longer-term resistance engaged under chronic milder stress, which might primarily reflect adaptability to stress.

We exposed aged worm cohorts to either acute oxidative-stress or heat-shock, seeking conditions that would kill 100% of worms within a few hours, relying on the timing of median survival to detect differences in stress resistance. Wild-type adult hermaphrodites exposed to 0.002% - 7% *tert*-butyl-hydroperoxide (*t*-BOOH) showed simple dose-dependency in oxidative stress resistance (**Fig. 1b**). Thermal stress, from 35°C - 42°C, also had a dose-dependent effect (**Fig. 1c**). To limit the influence of uptake mechanisms and stress adaptation on measured stress resistance, we used 7% *t*-BOOH and 42°C as acute stress challenges in subsequent studies, which are both characterized by a median time of death of 1.5 h.

We next tested these protocols on well-characterized longevity mutants, monitoring stress resistance throughout life, which the high-throughput capability of LFASS affords. The insulin/IGF-1 signalling (IIS) pathway is highly conserved from invertebrates to mammals, regulating growth, metabolism and ageing^19^. Mutations of the *daf-2* IIS receptor extend lifespan and increase stress resistance, while those of the downstream transcription factor DAF-16/FoxO shorten lifespan and cause stress hypersensitivity^20^. As expected, ageing cohorts of *daf-2* and *daf-16* mutants showed resistance and hypersensitivity, respectively, to acute stress (**Fig. 1e, 1f, Supplementary Table 1**). Against expectation, wild-type worms showed a *daf-16*-independent age increase in resistance to acute oxidative stress, which peaked at day 10 and reached a level of resistance comparable to that of stress resistant *daf-2* mutants (**Fig. 1e, Supplementary Tables 3, 4**). Because this effect persisted in animals with genetically or pharmacologically impaired feeding (**Supplementary Figure 2a, 2b**), and was supressed by specific mutations (**Fig. 2b-d**), it is unlikely to be due to an age-associated decrease in *t*-BOOH uptake. By contrast, wild-type animals exhibited a *daf-16*-dependent rise and fall in heat-shock resistance, peaking at day 6 (**Fig. 1f, Supplementary Table 2, 4**).

**Figure 2.**
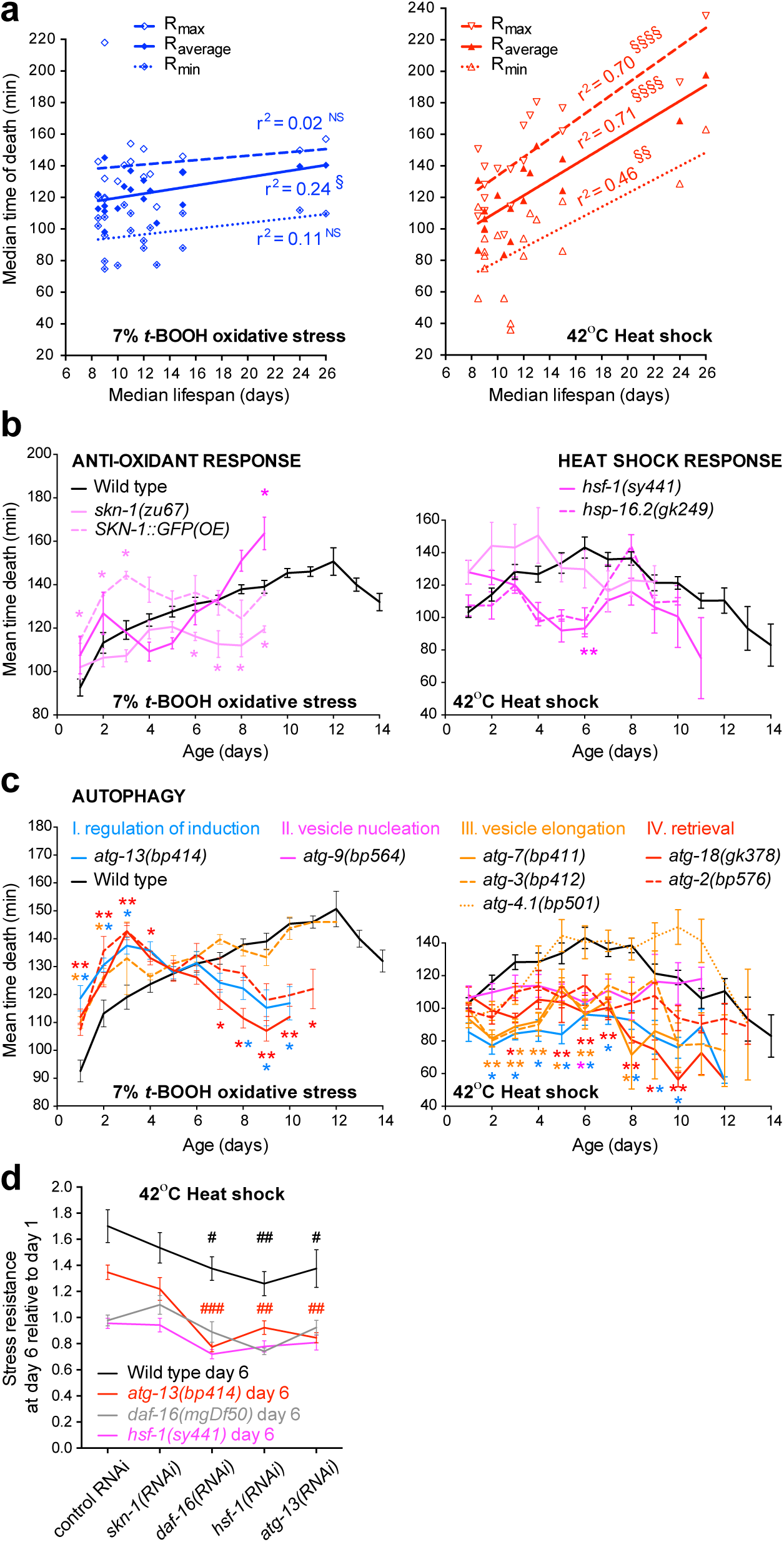
LFASS analysis of acute stress genetic determinants and correlation with longevity. (a) Minimum, mean and maximum heat-shock resistance strongly correlate with median lifespan, while correlation with acute oxidative-stress resistance is poor (§ *p*<0.05, §§ *p*<0.01, §§§§ *p*<0.0001). (b) *skn-1* is specifically required for the ageing-associated increase in acute oxidative stress resistance (left), while *hsf-1* is specifically required for ageing-associated increase in heat-shock resistance (right). (c) Autophagy is required for ageing-associated increase in both acute oxidative-stress (left) and heat-shock (right) resistance. Comparison with age-matched wild-type: * *p*<0.05 down to *p*<0001. (d) Adult-specific RNAi against *daf-16*, *hsf-1* and *atg-13* reduce age-associated heat-shock resistance. Comparison with control RNAi: # *p*<0.05, ## *p*<0.01, ### *p*<0.001. Error bars, SEM.

To explore the genetic basis of these phenomena, we screened selected ageing mutants. As manual analysis of high-throughput data generated by LFASS is very time consuming, we developed a software package to automatically extract median time of death from DF curves (**Supplementary Figure 3**, “http://www.ucl.ac.uk/iha/LFASS” www.ucl.ac.uk/iha/LFASS - updates available upon request). After querying the user for key assay parameters, fluorescence time-lapse data are automatically sorted, smoothened, fitted and median times of death logged into a data output table. The <5% unfitted curves can be re-analysed individually with user guidance. Automated analysis yielded near identical results to manual analysis (**Fig. 1g**), but in 1/100^th^ of the time.

Screening ageing mutants showed, again, major differences in the relationship between longevity and resistance to acute oxidative stress versus heat-shock (**Supplementary Figure 2**). Firstly, acute oxidative stress resistance correlated very poorly with lifespan across genotypes or treatments, while there was a strong positive correlation between heat-shock resistance and lifespan (**Fig. 2a, Supplementary Figure 4**). The *skn-1* (*NRF2*) antioxidant transcription factor^21^ was required for the age increase in oxidative stress resistance, and the *hsf-1* heat shock transcription factor^22^ for the age increase in heat-shock resistance (**Fig. 2b**). Interestingly, autophagy proved to be essential for both effects (**Fig. 2c**), in line with its known increase in activity during early adulthood^23^ and stress protective effects^23, 24^. All autophagy mutations tested suppressed ageing-associated resistance to heat-shock and oxidative stress (**Fig. 2c, Supplementary Table 1**), apart from *atg-4.1(bp501)* which lacks one of the two partially redundant *C. elegans* homologues of ATG4^25^. Consistent with these results, adult-specific RNAi targeted against *daf-16*, *hsf-1* and *atg-13* could all reduce ageing-associated heat-shock resistance in wild-type and *atg-13(bp414)* worms (**Fig. 2d**), but did not significantly affect acute oxidative stress resistance (**Supplementary Figure 5**). Thus, adult expression of *daf-16*, *hsf-1* and *atg-13* all contribute to the ageing-associated increase in heat-shock resistance but are dispensable for acute oxidative stress resistance.

While chronic stress resistance declines with age^13-15^, we find that acute stress resistance increases in early ageing. Moreover, the dynamics of heat-shock and acute oxidative stress resistance with ageing differ markedly. A hypothesis that reconciles these results is that older animals, being under “physiological stress”, have a higher initial stress-buffering capacity, but lack the capacity to adapt to sub-lethal stress. The earlier fall in heat-shock resistance compared to acute oxidative stress resistance is consistent with the previously observed early decline in proteostasis^13^. Importantly, our correlation analysis predicts that screens for heat-resistant rather than prooxidant-resistant variants are more likely to yield new determinants of ageing.

In summary, LFASS proved applicable to a range of nematode survival assays and its high-throughput capabilities enabled new insights about stress resistance and ageing to be obtained. It is unbiased, easily implemented and versatile, requiring no added reagents and is compatible with transgenic, frail or immobile worms. It does not require a strict sample size so that worm loading can be automated (e.g. using worm sorters, automatic dispensers). It is applicable to modern screening platforms employing transparent materials (e.g. microfluidic chips, multi-well plates) and is highly cost effective. It will be advantageous to accelerate anti-helminthic drug discovery, and studies of toxico-pharmacology, bacterial infection and ageing.

## Acknowledgments

We thank S. Bar-Nun (Tel-Aviv University) for useful discussion, M. Ezcurra, M. Piper, and G. Charras for comments on the manuscript, and T.K. Blackwell (Joslin Diabetes Center) and E. Schuster (University College London) for providing strains. Some strains were supplied by the *Caenorhabditis* Genetics Center at the University of Minnesota (funded by the National Institutes of Health - Office of Research Infrastructure Programs P40 OD010440). The European FP-7 IDEAL programme and a Welcome Trust Strategic Award to D. Gems funded this work.

## Methods

***C. elegans* strains and handling.** The following strains were used in this study: N2 (wild type); DA597, *phm-2(ad597) I*; DA2123, *adIs2122 [lgg-1p::GFP::lgg-1 + rol-6(su1006)]*; DR1567, *daf-2(m577) III*; DR1572, *daf-2(e1368) III*; EFS7, *daf-16(mgDf50) I*; *daf-2(e1370) III*; EU1, *skn-1(zu67) IV/nT1 [unc-?(n754) let-?] (IV;V)*; GA60, *eat-2(ad1116) II*; GA82, *daf-2(e1370) III*; GA91, *daf-16(mgDf50) I*; *daf-2(m577) III*; GA1001, *aak-2(ok524) X*; GR1307, *daf-16(mgDf50) I*; GS776, *unc-32(e189) lin-12(n676n930) III; unc-42(e270) sel-11(ar39) V*; GS807, *unc-32(e189) lin-12(n676n930) III; unc-42(e270) sel-11(ar39) V*; HZ1683, *him-5(e1490) V; atg-2(bp576) X*; HZ1684, *atg-3(bp412) IV; him-5(e1490) V*; HZ1685, *atg-4.1(bp501) I*; HZ1686, *bnIs1 I; atg-7(bp411) IV; him-5(e1490) V*; HZ1687, *atg-9(bp564) him-5(e1490) V*; HZ1688, *him-5(e1490) V; atg-2(bp576) X*; LD1004, *Ex001[SKN - 1B/C::GFP]*; PS3551, *hsf-1(sy441) I*; RE666, *ire-1(v33) II*; RB545, *pek-1(ok275) X*; RB938, *vha-12(ok821) X*; RB1021, *crt-1(ok948) V*; SJ17, *xbp-1(zc12) III; zcIs4 V*; SJ30, *ire-1(zc14) II; zcIs4 V*; SJ4005, *zcIs4[hsp-4::GFP] V*; SJ4100, *zcIs13[hsp-6::GFP]*; ST6, *eat-20(nc4) X*; VC475, *hsp-16.2(gk249) V*; VC893, *atg-18(gk378) V*; ZG31*, hif-1(ia4) V*. GA60, GA82, GA91, GA1001 were generated in the Gems lab. LD1004 was kindly provided by Keith Blackwell, and EFS7 by Eugene Schuster. All other strains used in this study were provided by the *Caenorhabditis* Genetics Center, University of Minnesota).

*C. elegans* strains were maintained at 15°C following standard culture conditions^26^, on NGM 60 mm-diameter agar plates seeded with *E. coli* OP50. Ageing worm cohorts were prepared as follows. Young adult hermaphrodites (30 per plate) were allowed to lay eggs for 24 hr. Several days later L4 animals were collected and transferred to NGM/OP50 plates containing 15 μM fluorodeoxyuridine (FUdR) (Sigma-Aldrich #F0503) to block egg production; or to NGM with 15 μM FUdR and 25 μg/mL carbenicillin (Sigma-Aldrich #C3416), and seeded with HT115 RNAi-producing bacteria, as described^27^, and in each case maintained at 25°C. L4 larvae were collected in this way daily for 2-3 weeks, and then adult hermaphrodites of each age were picked into multi-well plates and subjected to LFASS all on the same day.

**Time-lapse microscopy experiments.** 100-120 1 day old adult hermaphrodites were mounted in M9 on 2% agarose pads between slide and coverslip without anaesthetic unless otherwise stated. Imaging was performed through a DAPI filter set (Chroma technology Corp, USA) using a 2.5x objective on a Leica DMRXA2 microscope (Leica Biosystems Nussloch GmbH, Germany). Successive brightfield and DAPI images were acquired every 30s using the Volocity 6.3 software (Perkin Elmer, USA). For rapid killing experiments (**Supplemental Figure 1a**), a 70% solution of *tert*-butylhydroperoxide (Luperox TBH70X, Sigma-Aldrich #458139, Switzerland) was added volume to volume to the mounting M9 medium prior to coverslip apposition. The imaging protocol was started exactly 1 min later. For heat-killing experiments (**Supplemental Figure 1b**), a PE120 heating/cooling platform (Linkam Scientific Instruments Ltd, UK) was attached to the microscope stage. The heating protocol (ramping up to 42°C at 1.2°C per min) was initiated simultaneously with the time-lapse imaging.

**Time-lapse microscopy analysis.** We used the Volocity 6.3 Quantitation module to generate graphic representations (kymographs) of single worm traces from the 2.5x time-lapse imaging series. The time of death for each worm was deduced from the time of the intestinal blue fluorescence burst. Individual times of death during a single time-lapse were fitted into bins and count distributions plotted and fitted with a Gaussian curve using GraphPad Prism 6.0 software (GraphPad Software Inc., USA). Overall fluorescence for each time point was measured using the ImageJ-based open-source package Fiji (http://fiji.sc/Fiji), plotted and analyzed using GraphPad Prism 6.0.

**Plate-reader assays.** For oxidative stress and heat shock assays, we picked 16 worms into 60 μL M9 per well for 384-well plates, and 50 worms in 150 μL M9 for 96-well plates, together with a pellet of *E. coli* OP50 bacteria to prevent starvation. For infection assays, *Enterococcus faecalis* GH10 bacteria were streaked onto Brain Heart Infusion Kanamycin (BHIK) agar plates and used within a week, as described^28^. Liquid (BHI) *E. faecalis* cultures were grown for 3-5h at 37°C to saturation on the day. We then picked 100 worms per well into 50 μL M9 + 30 μL OP50 medium (for 384-well plates), and supplemented with 10 μL freshly saturated *E. faecalis* solution cooled to room temperature. A Tecan Infinite 200 plate-reader (Tecan Group Ltd., Switzerland) was pre-warmed at 25°C to match the temperature at which aged cohorts were raised and *E. faecalis* infections assays performed. Blue fluorescence (excitation: 360 nm / emission: 435 nm) was recorded for each well every 2min for 8h or every 5min for 4 days for stress and infection assays, respectively.

**Death fluorescence (DF) curve manual analysis.** Fluorescence time-lapse recording data for each well were normalised. The maximum was chosen where a significant peak of fluorescence was observed. Fluorescence values for the first 15-20 time points were often inaccurate, yielding local maxima and minima, due to worm thrashing in the wells, and were therefore omitted for the determination of the fluorescence minimum and maximum. After normalization, the time of half-maximum fluorescence was determined.

**DF curve automated analysis.** Matlab 2014b and 2015a versions were used to write and execute the LFASS software package. **Supplemental Figure 3** describes the approach. Detailed documentation (description of functions and variables) is provided together with the package (LFASS.zip) within the Readme_LFASS.txt file. Briefly, the program proceeds as follow.

(1) Matlab separates text and number matrices so that tags and values are stored in separate matrices. In .xlsx plate-reader files each row represents the fluorescence of a single well over time. The last column is the well identity/tag (attributes of the sample: age, genotype, drug treatment, bacterial type).

(2) For the fit to perform optimally and return median time of death in minutes, several assay parameters have to be informed by the user (see supplemental online methods). The time interval between two measurements of the same well allows for the results to be expressed in minutes. The noise threshold allows for discarding empty wells, and wells in which no fluorescence peak can be detected. The noise fluorescence threshold should be chosen above the fluorescence values measured in empty wells, and below the peak blue fluorescence value sample-containing wells. This threshold also depends on the number of worms per well, and all data treated in the same bulk analysis should have roughly the same number of worms per well. Max and min have to be identified for the normalisation of the data. Early fluorescence values can greatly fluctuate due to worm thrashing/swimming (in the absence of anaesthetic) associated with high or low fluorescence values that can exceed the relevant maximum and minimum. For this reason, the user must indicate in which time intervals min and max are to be expected. These intervals usually exclude the first 10-20 time points and have to include all the times of minimum or maximum fluorescence of all the data sets included in one analysis. Once the data are normalised, the sigmoid fit needs to be constrained within initial and final plateaux that match the min and the max values. Because of inaccuracies in measurements, to chose the best plateaux, they need to be fitted over several time points. To achieve this, a tolerance threshold is given for the min/max (i.e., 0.5/0.95). The fit function will then take into account time points around the min/max that are found within these tolerance thresholds (between 0 and 0.5, and between 0.95 and 1, respectively) and stop looking for additional points beyond. This will effectively define a fit region that encompasses the death-associated blue fluorescence burst, ignoring all other parts of the curve (see green dotted lines in **supplemental figure 3 (3b) and (b)** lower panels).

(3) To speed up computing, the program first excludes rows that do not need to be fitted such as parameter rows (date, time points, temperature with time, etc.) by keeping only rows with at least 6 consecutives numbers (non-data rows contain letters). It then uses the noise threshold defined earlier to exclude data rows that do not have values above this threshold, which eliminates most of the empty wells (3a). Inaccuracies in measurements are associated with noise spikes that can complicate fitting. To limit their influence, the data are smoothened twice using the “smooth” function (4 other smoothing options were compared and this performed best). Min and max are found, data are normalized, and a fit interval is found. In the region to fit, curves are typically sigmoidal in shape. Because we are only interested in extracting the time of half-maximum (corresponding to the median time of death, Fig.S1), the critical region to fit is around this time point, and the sigmoid fit performs optimally.

(4) As the analysis progresses, a 4-column wide .txt result table is filled. Column 1 contains all the tags, column 2 reports the half-maximum time inferred from unfitted normalised raw data, column 3 reports half-maximum times obtained with the bulk fitting analysis, and column 4 reports the updated values obtained from bulk fitting and user-guided analyses. Dataset names and parameters (temperature, duration of assay) are filled in the first rows of column 1.“0” fill empty cells from non-data rows. “1” fill cells in data rows that did not pass the user-defined noise threshold.“NaN” fills cells in data rows that could not be fitted.

(5) When the fit does not converge for a given row, the user can re-analyse it giving attribute values that differ form the bulk analysis and that are better suited for this specific row. Typically, this allows for recovery of the 5% exploitable low quality data that are excluded by the bulk analysis. The result table is updated after each re-analysis until the user stops.

(6) Post-processing is performed in Microsoft Excel for data sorting and basic row/columns operations. Then statistical analysis and graphical representations are processed in GraphPad Prism.

**Lifespan assays.** With the exception of RNAi experiments, all worm cohorts used in reported stress or lifespan assays were hermaphrodites maintained at 15°C on OP50-seeded NGM plates and switched at the L4 stage to OP50 plates supplemented with 15μM FUdR, and subsequently maintained at 25°C.

**Statistics.** For lifespan statistics we used the JMP 12.01 Pro software package from SAS (USA). Lifespans were compared using the non-parametric log rank test. Unless otherwise stated, all other statistics were performed using Prism 6.0 from GraphPad Software Inc. (USA). Stress resistance differences with age and across genotypes were assessed by two-way ANNOVA with a post-hoc Dunnet’s test. *p* values reported in supplementary tables are adjusted for multiple comparisons.

---

**Supplemetary Table 1.**
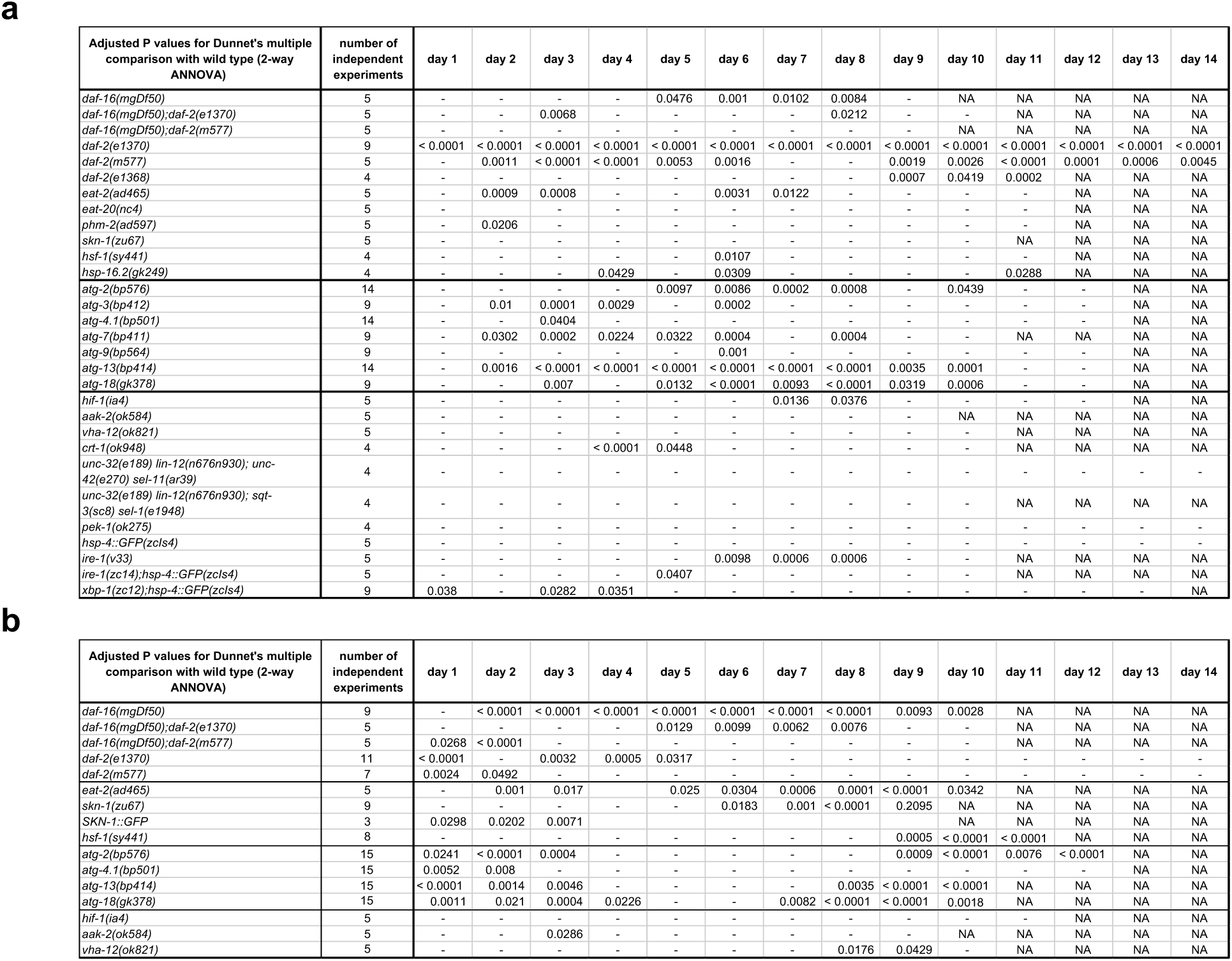
Supplemetary Table 1.

**Supplemetary Table 2.**
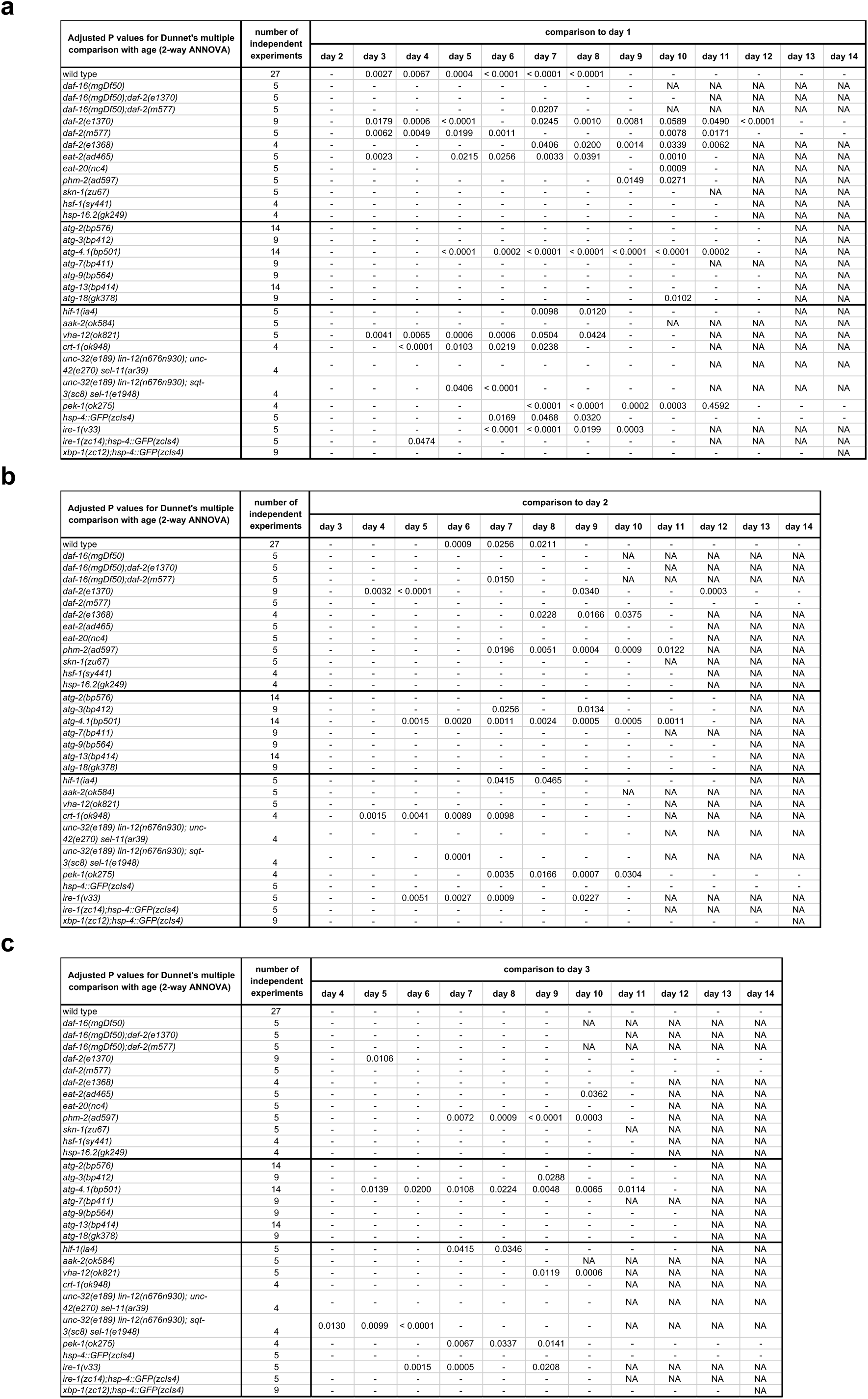
Supplemetary Table 2.

**Supplemetary Table 3.**
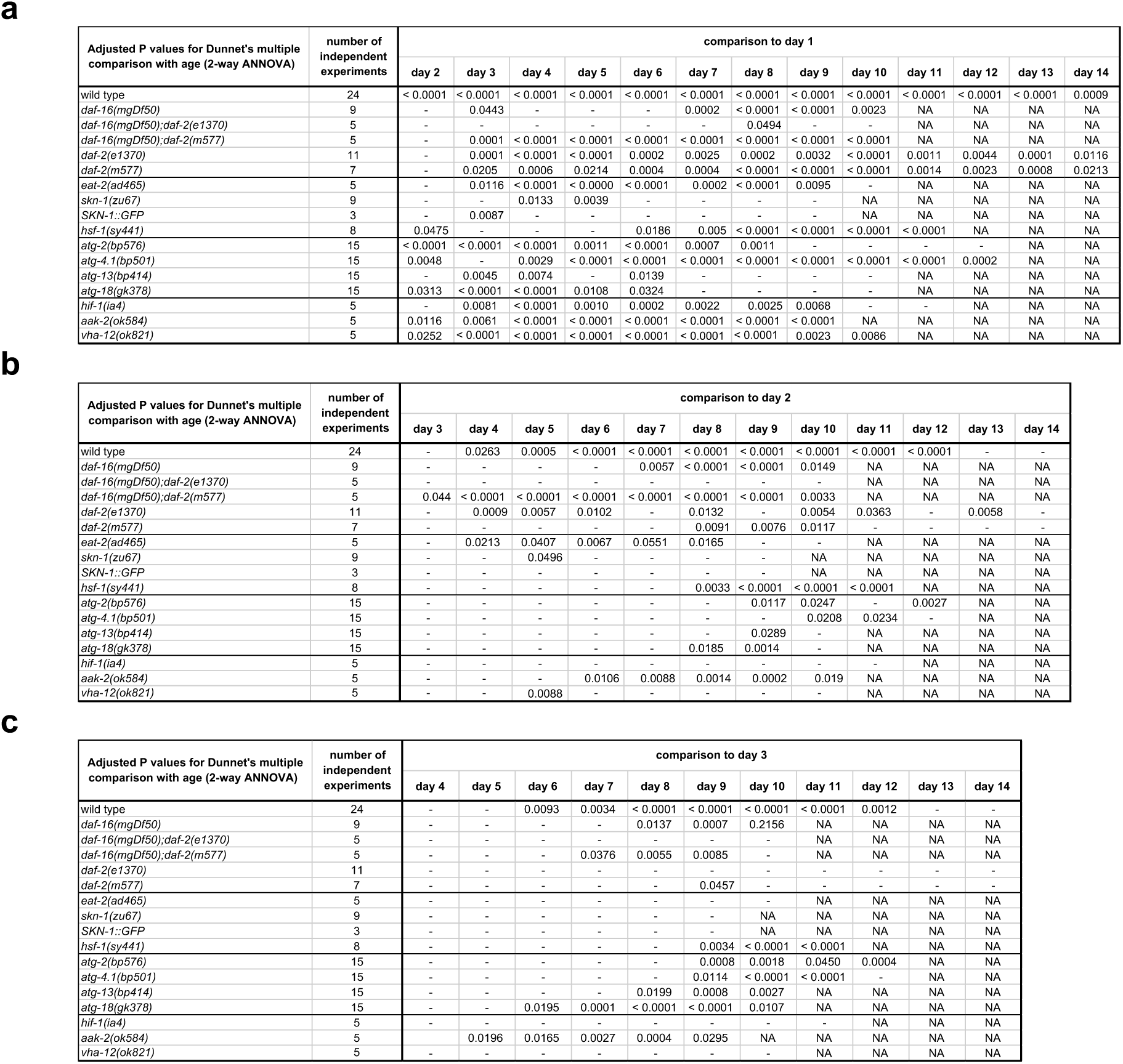
Supplemetary Table 3.

**Supplemetary Table 4.**
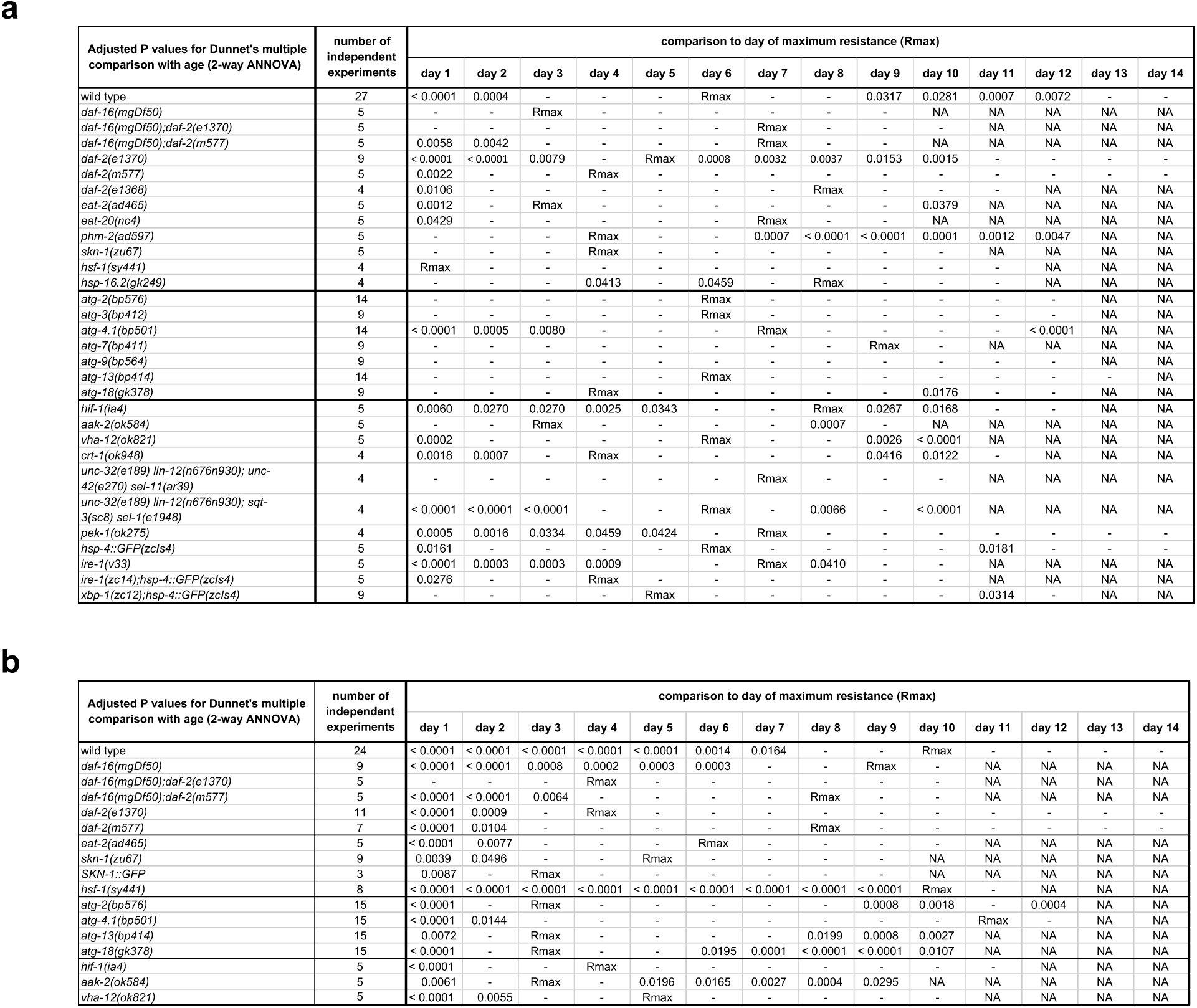
Supplemetary Table 4.

Supplementary legends

**Supplemental Figure 1.**
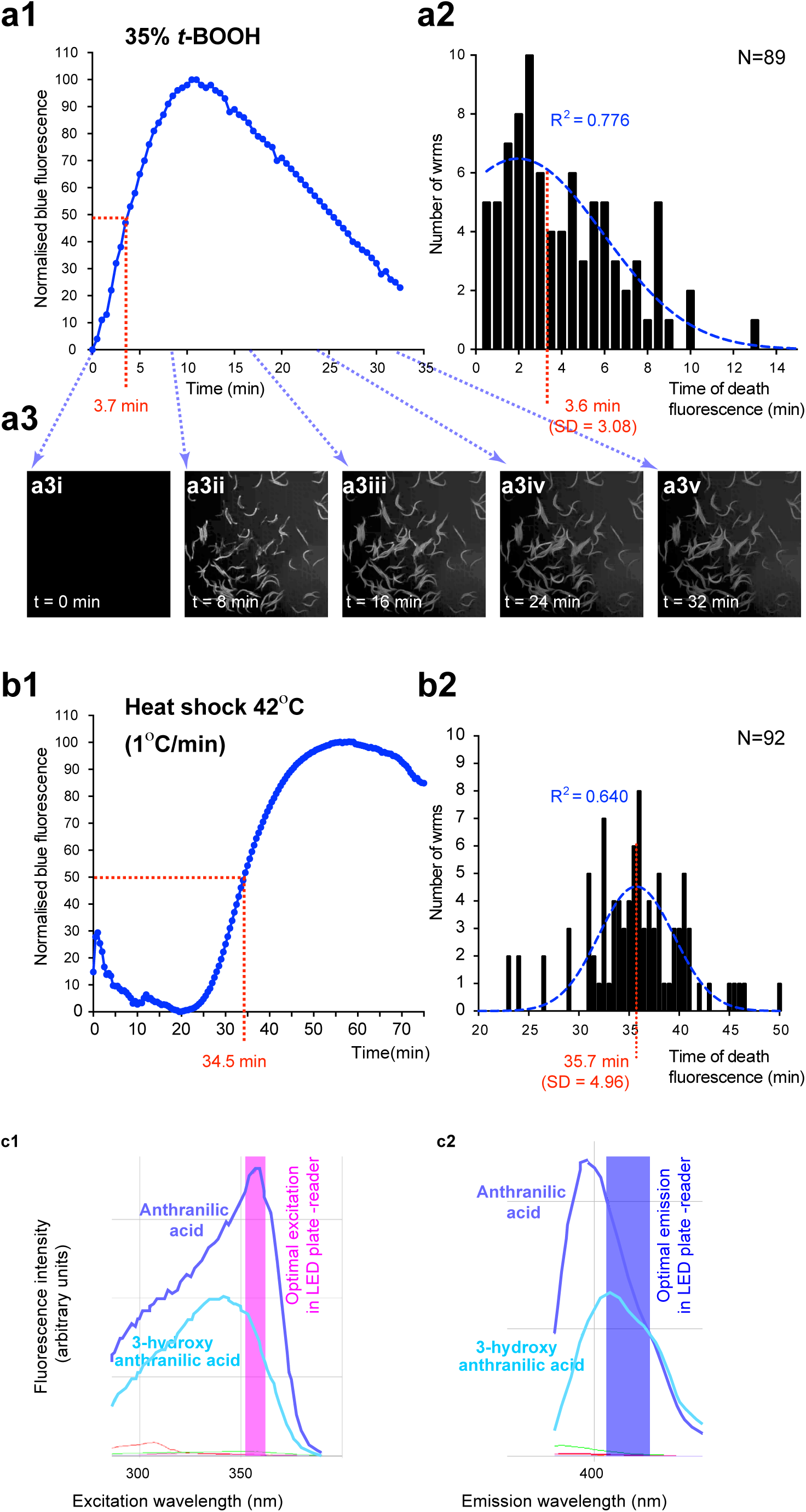
Using population death fluorescence to determine median time of death. Time-lapse microscopy (exemplified by panels a3i-a3v) of worm population submitted to fast (a) and slower (b) killing assays reveal that the time of half maximum death fluorescence (a1, b1, red dotted line) corresponds to the median time of death (a2, b2, red dotted line). Optimal excitation (c1) and emission (c2) windows for anthranilate-dependent death fluorescence in 384 well plates read by a LED plate-reader, matched with excitation and emission spectra for two commercially available anthranilate compounds.

**Supplemental Figure 2.**
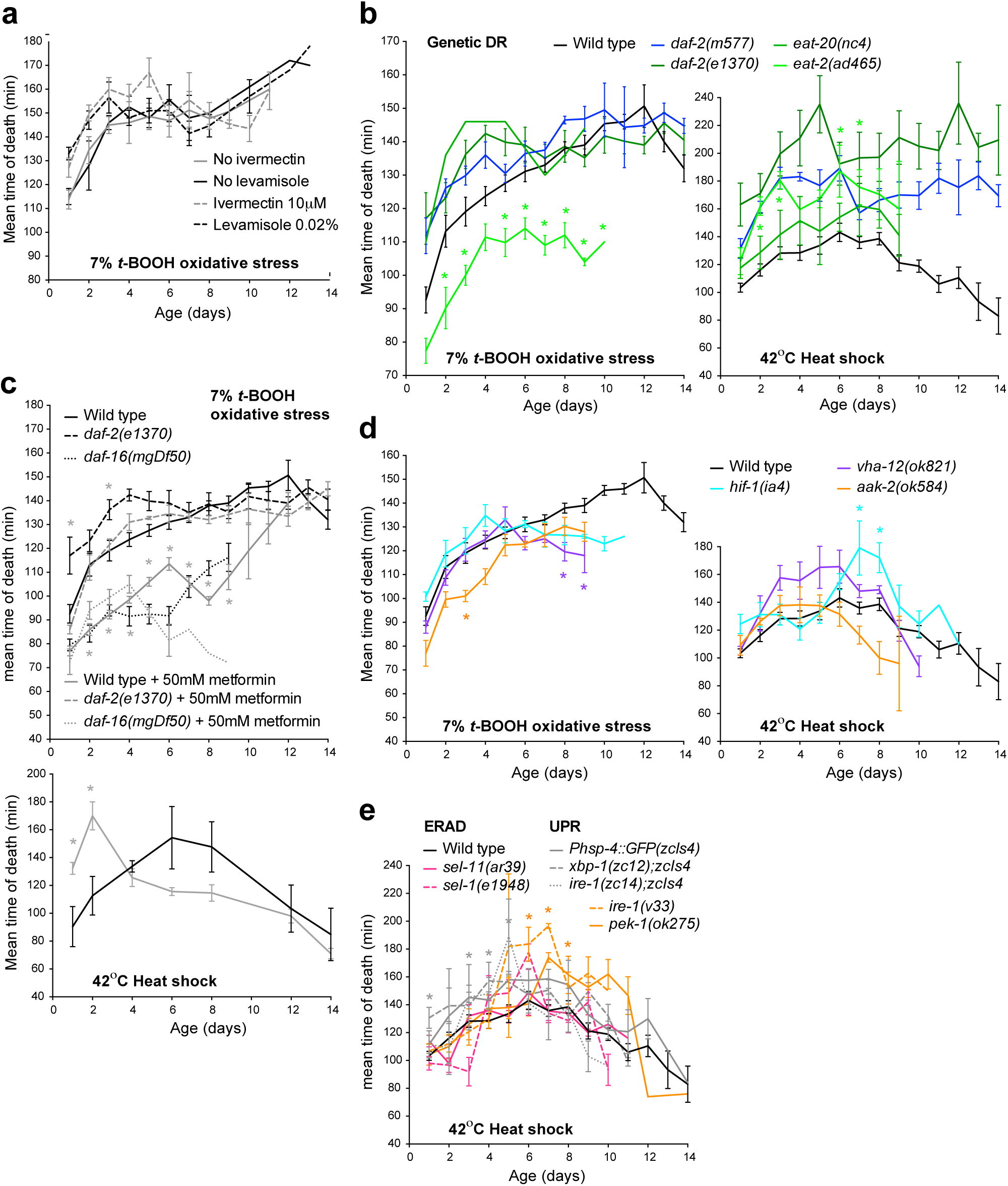
Effects of lifespan-affecting mutations and drug treatments on the age increase in acute stress resistance. (a) Inhibition of pharyngeal pumping by ivermectin or levamisole does not suppress the age increase in oxidative stress resistance. b) Slower pumping rate and associated genetically-induced dietary restriction (DR) do not correlate with acute oxidative-stress resistance dynamics (left) but correlates with heat-shock resistance dynamics: *eat-20(nc4)* is a mild pumping-defective mutant, *eat-2(ad465)* is a strong pumping/genetic DR mutant, *daf-2(e1370)* pumps more slowly than *daf-2(m577)* and so combines a genetic DR phenotype with reduced IIS. (c) The lifespan extending drug metformin sensitizes *daf-2* and wild type worms to acute oxidative stress but confers increased heat-shock resistance to young wild-type adults. (d) AMP-activated protein kinase alpha subunit (*aak-2*), vacuolar-H^+^-ATPase subunit 12 (*vha-12*) and hypoxia-induced factor 1 (*hif-1*) differentially affect age increases in acute oxidative-stress and heat-shock resistance. (e) The endoplasmic reticulum-associated protein degradation (ERAD) and unfolded protein response pathways are dispensable for the age increase in heat stress resistance. Error bars, SEM. Comparison with age-matched wild-type: * *p*<0.05 down to *p*<0001.

**Supplemental Figure 3.**
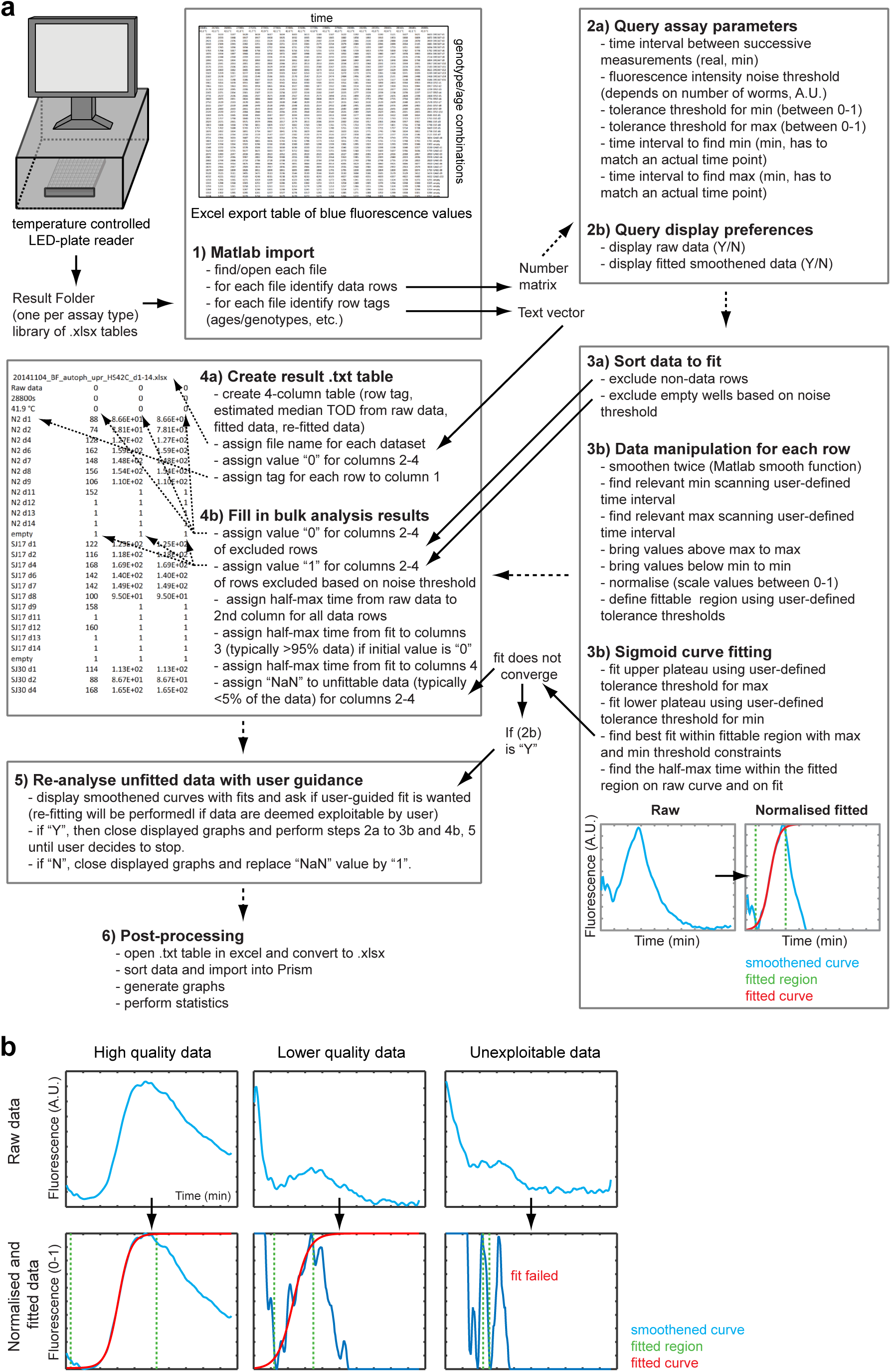
Diagram of the automated curve analysis package.a) Sequence of operations performed to automatically extract median time of death from raw time-lapse fluorescence data acquired by the LED-plate reader. (b) Examples of typical fluorescence curves and how the automated bulk analysis fits them. For bulk analysis, the same assays parameters are given for all data. The fitted region (green) adapts to the data, allowing for a good fit (red) to be found even on lower quality data (middle panels). Note that data deemed non-exploitable by visual inspection are not fitted (right panels) preventing erroneous data from filling the result table.

**Supplemental Figure 4.**
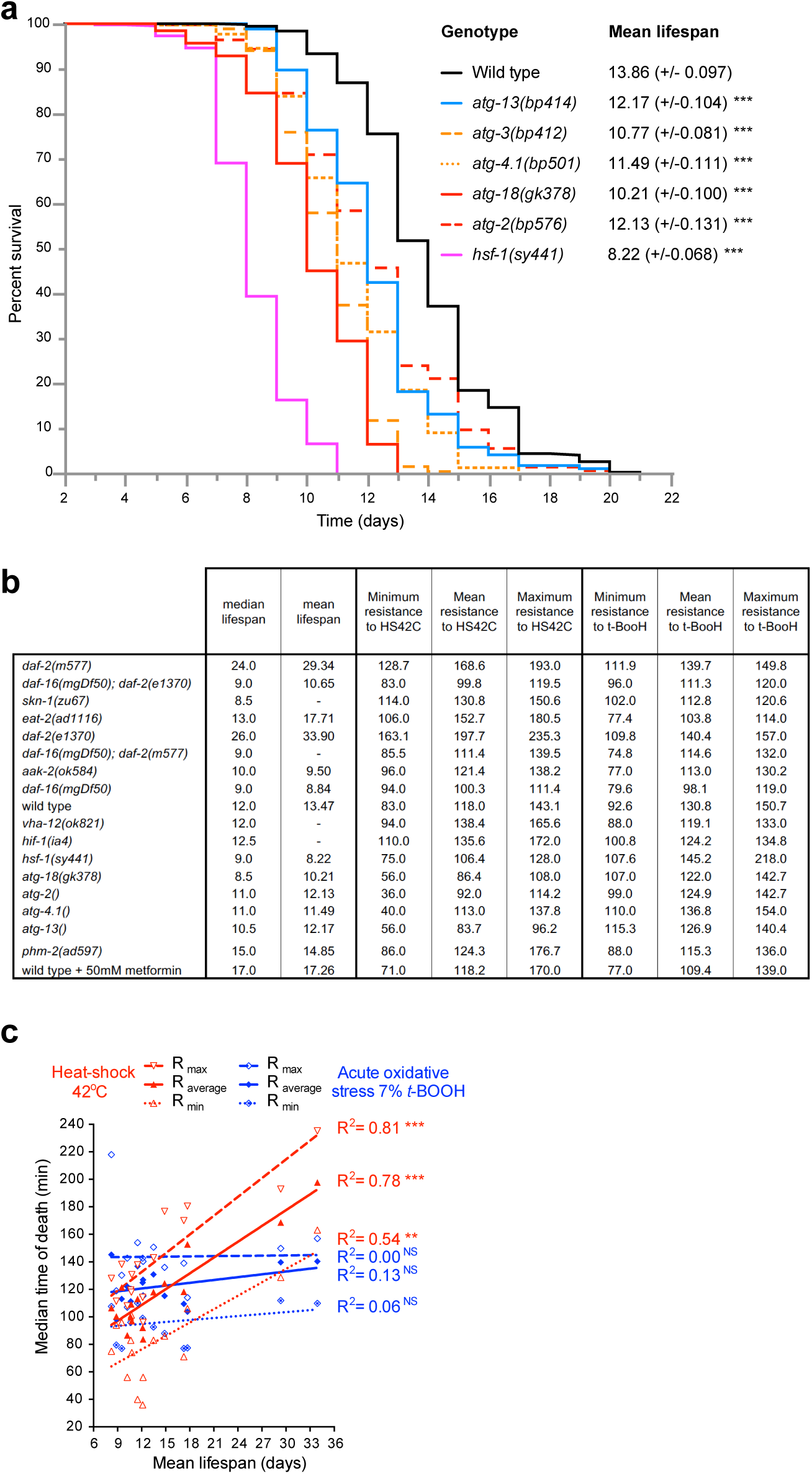
Correlations between acute stress resistance and lifespan.(a) Average lifespans of 4 independent experiments for the autophagy mutants used in the correlation analyses. (b) Table of median (matching estimates for the LFASS stress assays performed) and mean lifespan values (combining experiments performed in our lab and in the Cabreiro lab using the same growth conditions^29-31^, see Online Methods) used for the correlation plots in Figure 2a and Supplemental Figure 4c. Resistance levels are expressed in min, corresponding to the median time of death in LFASS assays. (c) Correlation plots between mean lifespan and acute oxidative-stress. Linear regressions are characterized by their correlation coefficient R.

**Supplemental Figure 5.**
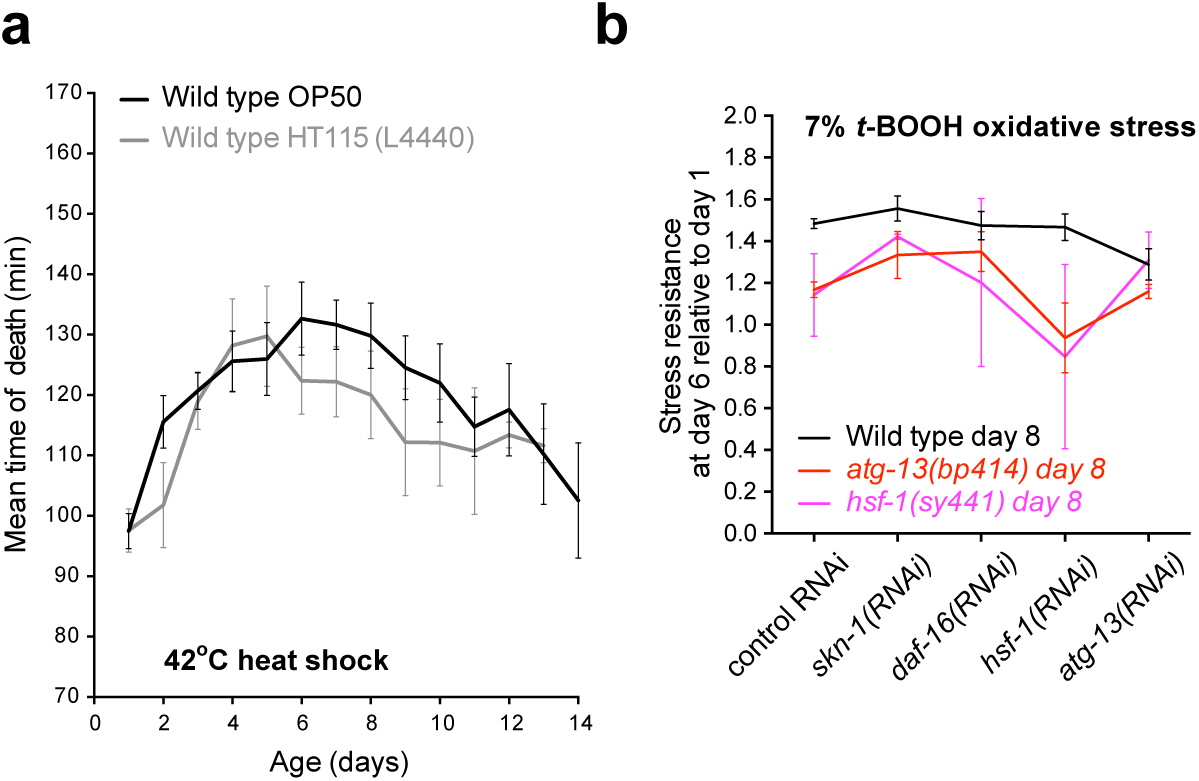
RNAi by feeding experiments.(a) *E. coli* strain HT115 diet does not significantly affect age increase in heat stress resistance. (b) Adult-specific

